# Obesogenic diet exposure alters uterine natural killer cell biology and impairs vasculature remodeling in mice

**DOI:** 10.1101/275503

**Authors:** Jennet Baltayeva, Chaini Konwar, Barbara Castellana, Danielle L. Mara, Julian K. Christians, Alexander G. Beristain

**Affiliations:** British Columbia Children’s Hospital Research Institute, Vancouver, Canada; Department of Obstetrics & Gynecology, The University of British Columbia, Vancouver,Canada; Department of Medical Genetics, The University of British Columbia, Vancouver, Canada; Department of Biological Sciences, Simon Fraser University, Burnaby, Canada

**Author notes:** To whom correspondence should be addressed: Alexander G. Beristain, British Columbia Children’s Hospital Research Institute, Department of Obstetrics & Gynecology, The University of British Columbia, Vancouver, British Columbia, Canada. V5Z 4H4.

**Keywords:** Pregnancy, natural killer cell, uterus, decidua, placenta, diet-induced obesity, spiral artery remodeling, high-fat/high-sucrose diet, gene expression, gene microarray

## Abstract

Pre-pregnancy obesity associates with adverse reproductive outcomes that impact maternal and fetal health. While obesity-driven mechanisms linked to poor pregnancy outcomes remain unclear, it is possible that obesity affects local uterine immune cells. Uterine immune cells, particularly uterine natural killer cells (uNK), play central roles in orchestrating developmental events in pregnancy. However, the effect of obesity on uNK biology is poorly understood. Using a high fat/high sugar diet (HFD) obesogenic mouse model, we set out to examine the effects of maternal obesity on uNK composition and establishment of the maternal-fetal interface. HFD exposure resulted in weight gain-dependent increases in systemic inflammation and rates of fetal resorption. Within HFD mice, both weight gain-dependent (diet-induced obese; DIO) and -independent (DIO-resistant; DIO-R) effects on natural cytotoxicity receptor-1 frequency and uNK activity were observed. Importantly, HFD-associated changes in uNK coincided with impairments in uterine artery remodeling in mid but not late pregnancy. Comparison of uNK mRNA transcripts in control diet mice to gene signatures in DIO or DIO-R identified differentially expressed genes that play roles in promoting activity/cytotoxicity and impeding vascular biology. Together, this work provides new insight into how obesity, and particularly how diet-induced weight-gain, may impact uNK-processes important in pregnancy.

## INTRODUCTION

Obesity affects more than half a billion adults worldwide and has become a global epidemic^1^. Consistent with trends in the general population, rates of obesity among women of childbearing age are increasing^2–4^. Pre-pregnancy obesity is recognized as a high-risk state because of its association with multiple adverse obstetric and perinatal outcomes including miscarriage, preeclampsia, preterm birth, and intrauterine fetal demise^5,6^. Providing the interface between the mother and developing baby, the conceptus-derived placenta plays an essential role in the transfer of nutrients and gases between maternal and fetal circulation. Importantly, placenta development and fetal health is impaired in obese women^7,8^. In rodent models of obesity, obesogenic diets promote placental dysfunction, defined in part by defects in utero-placental vascularization and altered trophoblast maturation^9–11^. However, the cellular and molecular processes affected by obesity that underlie detrimental obesity-related pregnancy outcomes are poorly understood.

Optimal placental function and overall pregnancy success is largely dependent upon tightly regulated angiogenic and vascular remodeling programs that take place within the maternal-fetal interface^12,13^. Early stages of uterine blood vessel growth, branching, pruning, and remodeling are controlled by diverse populations of leukocytes that reside within the decidual microenvironment^14,15^. In early human and mouse pregnancy, uterine natural killer cells (uNK) constitute the most abundant immune cell subtype within the uterine mucosa^16–18^. Genetic studies in mice show that uNK are major contributors to uterine vascular growth and remodeling, where these uNK-related processes are mediated in part by the production of pro-angiogenic factors (i.e. vascular endothelial growth factor; VEGF) and tissue-remodeling cytokines (i.e. tumor necrosis factor-α; TNF-α, type-II interferon)^19–21^.

Obesity associates with adipose tissue immune cell infiltration, including macrophages, T and B lymphocytes, mast cells, and neutrophils^22–24^, and along with these changes, obesogenic environments alter leukocyte activation by potentiating the secretion of pro-inflammatory cytokines that have effects within both local and distal tissues^25^. In women, previous work shows that uNK frequencies decrease in obesity^26^. Importantly, these cellular changes associate with increased rates of uNK degranulation, changes in natural killer receptor expression, and decreased production of pro-angiogenic VEGF-A^27^. While these findings provide insight into how maternal obesity affects uNK processes important in pregnancy, due to the heterogeneous nature of obesity within human populations, the need for a clearer understanding of how excess adiposity impacts uNK biology and how these changes affect pregnancy outcome is still needed.

In this study, we established a diet-based mouse model to investigate the impact of obesity on pregnancy outcome, uNK function, and placental development in early-mid pregnancy. Mice subjected to a chronic high-fat/high-sugar (HFD) diet showed increased levels of systemic inflammation and increased rates of fetal resorption. Notably, uNK composition within HFD mice was different than in mice fed a control diet, where alterations in state of uNK activity and receptor expression were observed. Notably, these uNK-related differences associated with uterine artery remodeling impairments within HFD mice. While diet exposure did not result in robust global gene expression changes in uNK, specific genes important in vascular biology, inflammation/cytotoxic activation, and metabolism were altered by HFD exposure. Together, our work provides novel insight into obesity-induced changes in uNK function. These findings also establish insight into immunological mechanisms impacted by maternal obesity that likely lead to alterations in utero-placental development in early-mid pregnancy which in turn may contribute to the development of obesity-linked pregnancy disorders.

## RESULTS

### Diet-induced weight gain drives systemic inflammation and potentiates fetal resorption

To investigate the effects of maternal obesity on uNK biology and placental development, female C57BL/6J mice were fed either a control low-fat/no sucrose diet (LFD) or a HFD over a 13-week period prior to mating; exposure to these diets was also maintained during pregnancy (Figure 1A). As expected, mice fed a HFD showed an increase in body weight (Figure 1B). However, there was considerable heterogeneity in endpoint body weight within the HFD cohort, where weight gain rates in ~37% of HFD mice (7/19) were indistinguishable from LFD controls (Figures 1C, 1D). This observation led us to sub-categorize mice within the HFD group as either weight-gainers (diet-induced obese mice; DIO) or as weight-gain resistant mice (DIO-resistant; DIO-R) (Figure 1C). Subcategorization of DIO and DIO-R mice was established following the example of Enriori *et al*^28^, where mice with endpoint weights > 3 standard deviations to the mean LFD weight were categorized as DIO. Consistent with previous findings showing association of HFD-induced weight gain and increased systemic inflammation^29,30^, levels of pro-inflammatory TNFα in blood serum were higher in DIO mice than in LFD controls at gestational day (Gd)10.5 (Figure 1E). Notably, a statistical difference was not observed in TNFα serum levels between DIO-R and LFD mice, suggesting that the increase in TNFα is due to increased adiposity and not exposure to HFD *per se* (Figure 1E).

**Figure 1.**
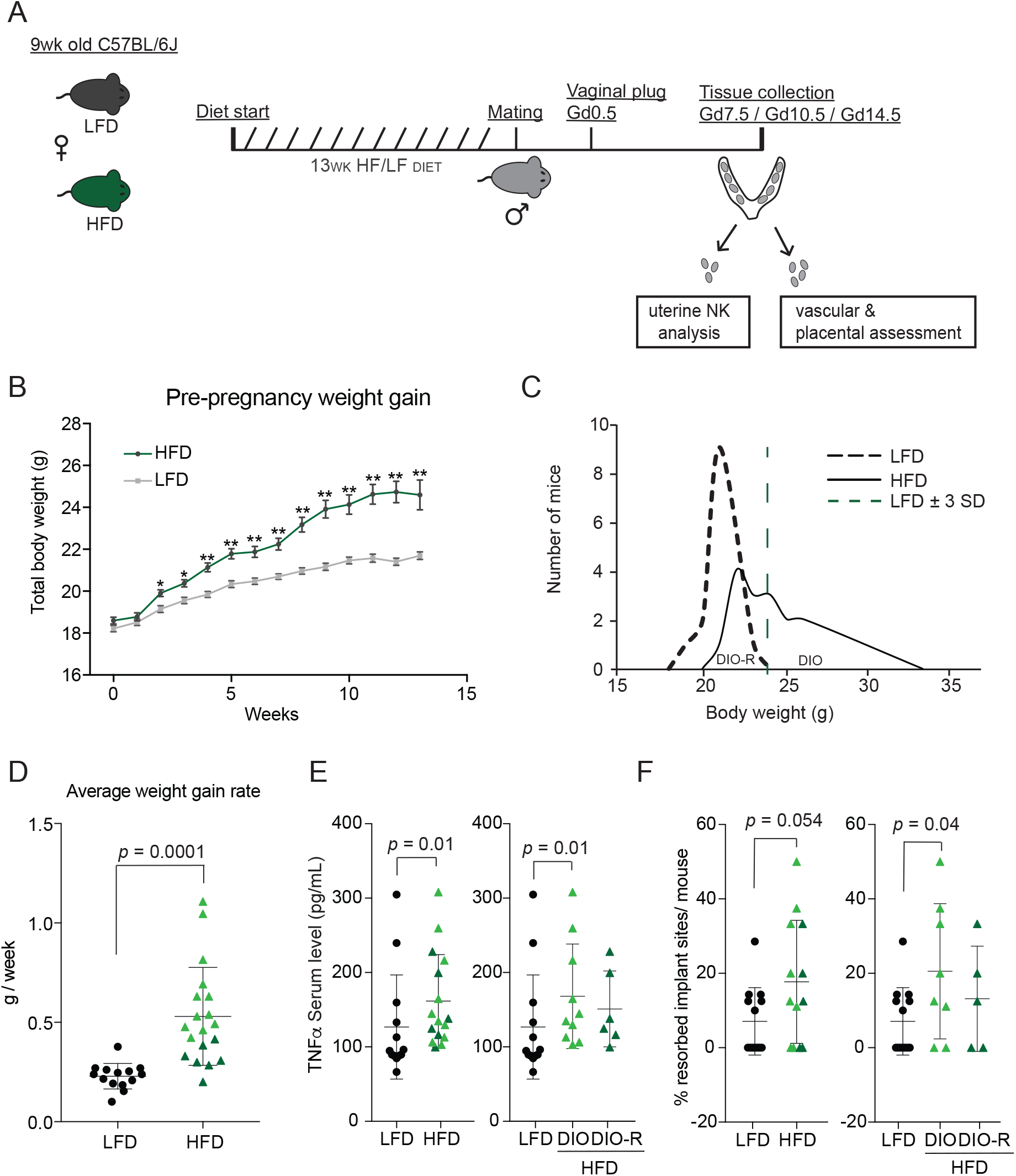
Establishment of a mouse model of diet-induced maternal obesity. **A)** Schematic diagram depicting the experimental workflow with low-fat, no sucrose (LFD) or high-fat, high sucrose (HFD) diet regime, breeding strategy, pregnancy detection at gestational day (Gd) 0.5, and gestational endpoints (Gd 7.5/10.5/14.5). Shown are uterine horns with multiple conception sites comprised of utero-placental tissues that were allocated for downstream cellular analyses or histology. **B)** Body weight gain prior to pregnancy of mice fed either HFD (black bars; n=20) or a matched LFD (grey bars; n=20). Data represented as mean ± SEM. *p<0.05; **p<0.005. **C)** Body weight distribution of LFD- and HFD-exposed mice following the 13-week diet regime. Red dashed line represents the mean LFD weight + 3 SD that was used to subcategorize HFD mice into diet-induced obese (DIO) and diet-induced obese resistant (DIO-R) groups. **D)** Average weekly weight gain rate (LFD: n=14, HFD: n=19). Data represented as mean ± SD. *P*-values from Student’s t-test. **E)** Maternal serum levels of TNF-α at Gd10.5 in LFD (n=13), HFD (n=16), DIO (n=10), and DIO-R (n=6) mice. Data represented as mean ± SD. *P*-values from Mann-Whitney test. **F)** Proportion of resorbed implantation sites per mouse observed at Gd14.5 (LFD: n=13, HFD: n=13); Data represented as mean ± SD. *P*-values from Student’s t-test.

Neither HFD exposure nor increased weight gain led to differences in placental or fetal measurements, although a trend for DIO-R placentas to weigh less than control placentas was observed (Supplemental Figure 1A, B). However, HFD exposure led to moderate, but significant increases in the fetal resorption rate compared to control mice (18% vs. 5%), and this effect was primarily driven by the DIO subgroup (Figure 1F). Together, our mouse model of maternal obesity shows that HFD-induced weight gain in pregnancy correlates with increased systemic inflammation and impaired fetal viability.

### High-fat diet exposure does not alter uNK frequencies in early pregnancy

Uterine NK in mice are defined as “tissue-resident” Dolichos biflorus agglutinin (DBA)^+^CD49a^+^DX5^+/-^ (tr-uNK) or “conventional” DBA^−^CD49a^−^DX5^+^ (c-uNK), and these distinct subtypes play specific roles in uterine neo-angiogenesis (i.e. tr-uNK) and artery remodeling (i.e. c-uNK)^31^. In mid pregnancy, tr-uNK are the dominant uNK subtype comprising roughly 80% of all uNK, while c-uNK make up a smaller proportion^32^. To this end, we first investigated whether HFD exposure alters the number of tr-uNK in early pregnancy by histologically examining DBA^+^ cells at Gd7.5 and Gd10.5 of pregnancy; at Gd7.5, uNK begin to expand in number and this expansion peaks by Gd10.5, at which point numbers subsequently decline^33^. While DBA^+^ uNK in both HFD and LFD mucosa underwent normal gestational age-related increases in numbers, uNK numbers were not affected by diet (Figure 2A).

**Figure 2.**
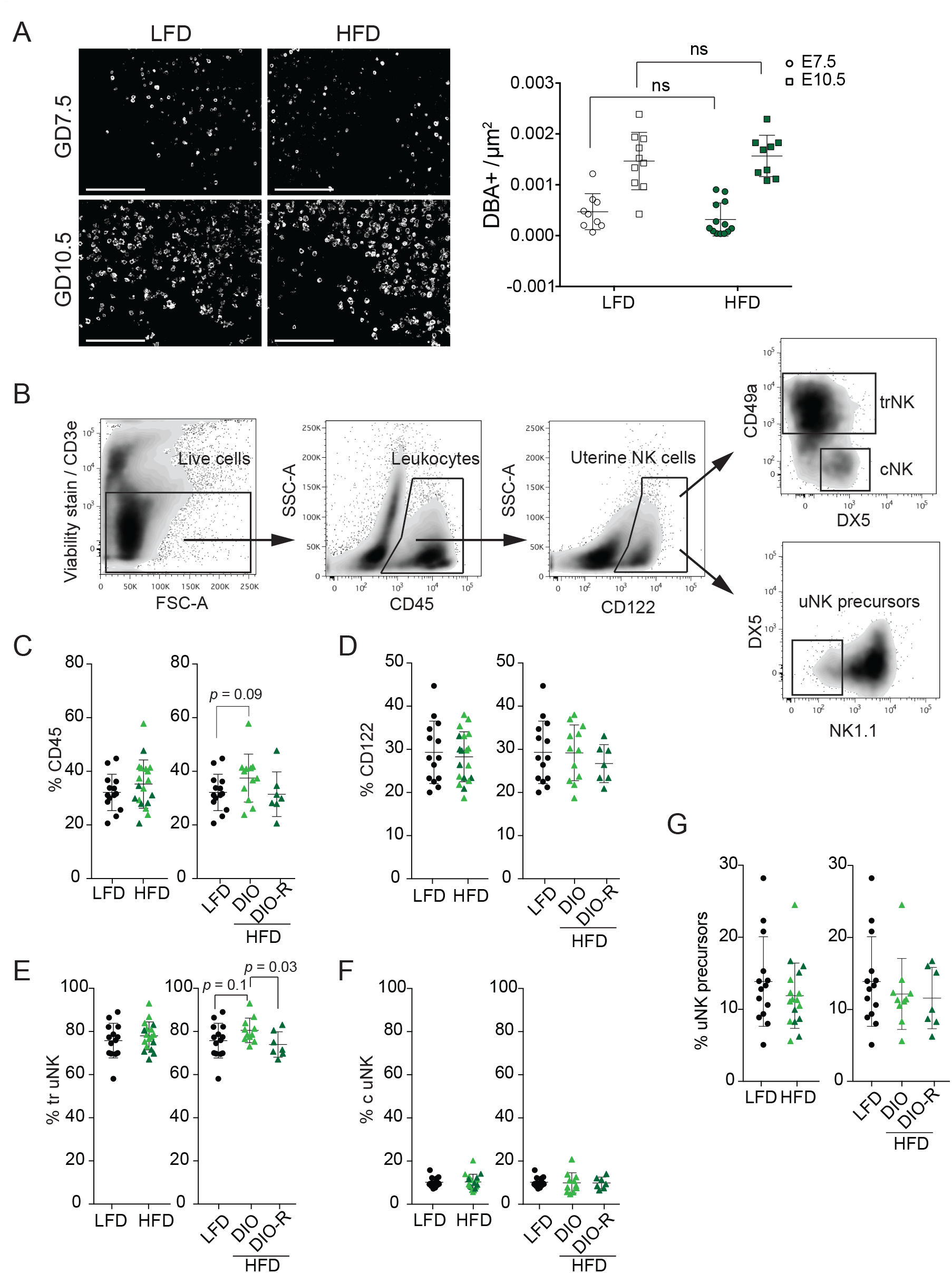
HFD exposure does not result in alterations of total uterine leukocytes or subsets of uNK cells. **A)** Representative DBA immunofluorescence images of implantation sites at Gd7.5 (LFD: n=9; HFD: n=13) and Gd10.5 (LFD: n=10; HFD: n=9) and scatter plots showing number of dolichos biflorus agglutinin (DBA)+ uterine NK cells. Scale bar = 200μm. **B)** Flow cytometry gating strategy identification and quantification of uterine immune cell populations, including total uNK (CD3e-/CD45+/CD122+), tissue-resident (tr) (CD122+/CD49a+), conventional (c) (CD122+/DX5+), and uNK precursors (CD122+/DX5-/NK1.1-). Representative scatter plots of **C)** total leukocyte (CD45+), **D)** uterine NK, **E)** tr-uNK, **F)** c-uNK, and **G)** uNK precursors in LFD (n=14), HFD (n=18), DIO (n=11), and DIO-R (n=7) mice. Data represented as mean ± SD; *P*-values from unpaired Student’s t-test for panels A, C-G.

Following these histological assessments, proportions of total leukocytes (CD45^+^) and uNK (CD3^-^CD122^+^) were examined by flow cytometry in Gd10.5 implantation sites, as were specific truNK, c-uNK, and uNK precursor cell sub-populations (Figure 2B). HFD exposure showed a modest, but non-significant effect on total CD45^+^ immune cell proportions, where an increasing trend in CD45^+^ cell frequency was observed only within the DIO subgroup (Figure 2C). Similarly, total uNK frequencies were not impacted by HFD or diet-induced weight gain (Figure 2D). Consistent with this, HFD did not exert any measurable differences in proportions of tr-uNK, although an increase in tr-uNK frequency was observed within DIO compared to DIO-R mice; a similar though non-significant relationship (*P* = 0.1) between tr-uNK proportions in DIO and LFD control mice was also observed (Figures 2E). Conventional-uNK made up a smaller proportion (~ 10-20%) of uNK compared to the dominant tr-uNK population, where frequencies were also not altered by diet or diet-induced weight gain (Figure 2F). Lastly, the effect of HFD on uNK precursors (CD122^+^/NK1.1^-^/TDX5^-^)^34^ was measured, and consistent with the above findings, HFD exposure did not affect precursor frequencies (Figure 2G).

### Proportions of NCR1^+^ uNK increase in response to high-fat diet

Natural cytotoxicity receptor 1 (NCR1) is essential for uNK maturation and function, and plays critical roles in promoting uterine blood vessel remodeling in mice^35,36^. Thus, to further elucidate the effects of maternal HFD on uNK biology, NCR1 uNK proportions and expression levels were examined by flow cytometry. While HFD did not impact cell surface levels of NCR1 expression (MFI), mice exposed to HFD did show increased frequencies of NCR1^+^ uNK (Figures 3A, B, C). Further analysis revealed that the increase in NCR1^+^ uNK in HFD mice is primarily driven by the DIO-R subgroup; DIO mice showed a modest but non-significant increase in NCR1^+^ uNK proportions (*P* = 0.09) (Figure 3B). These findings suggest that obesogenic diet exposure, and to a lesser extent diet-associated weight gain, manifest in subtle changes in uNK composition defined by NCR1 expression. Given the importance of NCR1 in vascular processes in pregnancy, diet-induced differences in NCR1 may translate to alterations in uterine artery remodeling or cytotoxicity.

**Figure 3.**
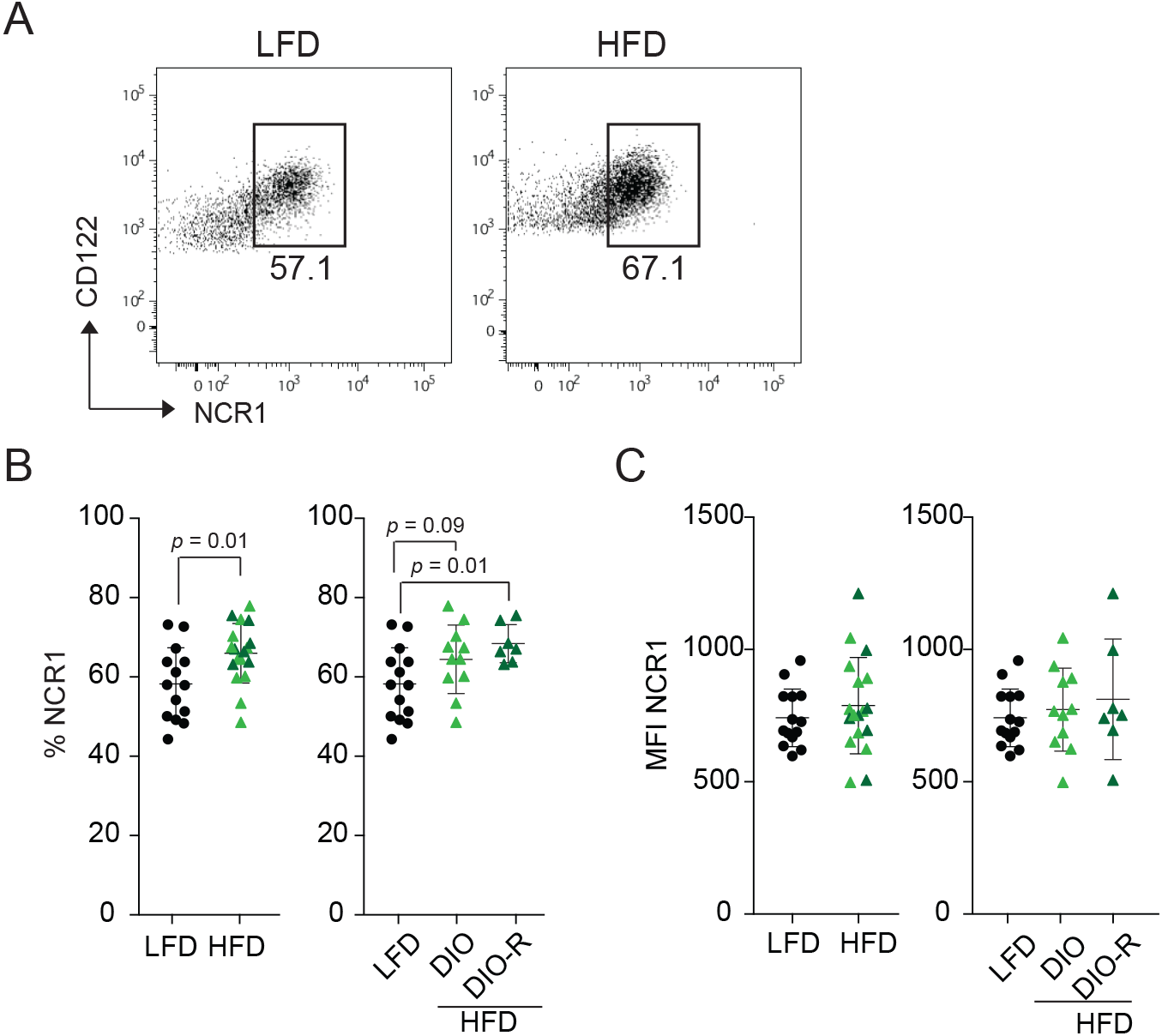
HFD and diet-induced weight gain alter frequencies of NCR1+ uNK cells. **A)** Representative flow cytometry plots of NCR1+ uNK proportions. Scatter plots show **B)** proportion of NCR1+ uNK and **C)** NCR1 median fluorescent intensities (MFI) readings within uNK of LFD (n=14) HFD (n=18), DIO (n=11), and DIO-R (n=7) mice.

### HFD exposure drives uNK activity

Since NCR1 associates with the promotion of uNK maturation and activation, we next set out to examine if HFD exposure alters uNK activity^36^. To do this, two approaches were used: Immunofluorescence microscopy facilitated measurement of uNK cell diameter/size as a readout of maturity^36^, and flow cytometry analysis measured cell surface expression of the early activation marker CD69^37^. Because NCR1 is expressed predominantly by DBA^+^ cells at Gd10.5 ^36^, focus was given to measuring DBA^+^ cell diameters within histological sections of utero-placental tissues. While cell diameters of DBA^+^ uNK were not shown to be different between HFD and control LFD mice (Supplemental Figure 1C), CD69 expression levels (MFI) were significantly higher in both total (CD3^-^ CD122^+^) and NCR1^+^ uNK populations as a result of HFD exposure (Figures 4A, B, C). In line with our findings that NCR1 proportions are higher within the DIO-R sub-group than in control LFD or DIO mice, the increase in HFD-induced CD69 expression was primarily attributed to uNK within the DIOR, and not the DIO subgroup, the latter of which showed greater heterogeneity in CD69 expression (Figures 4B, C). Overall these findings suggest that HFD exposure, independent of weight gain, instructs an altered state of uNK activity.

**Figure 4.**
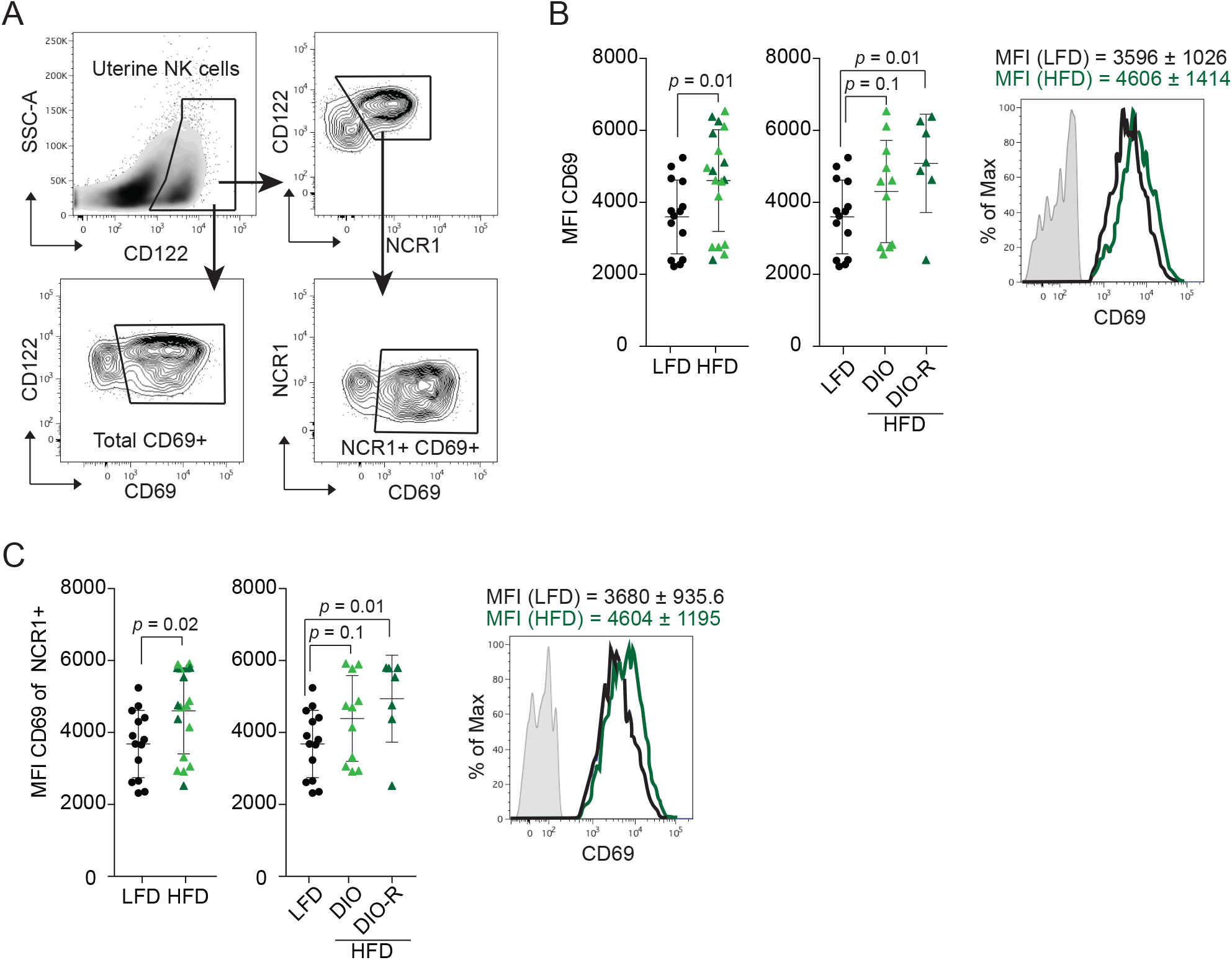
High-fat diet and diet-induced weight-gain potentiates uNK activity. **A)** Representative flow cytometry plots and gating strategy for analyzing proportions of CD69-expressing uNK and NCR1+ uNK. **B)** Representative MFI histograms and scatter plots showing CD69 expression intensity (MFI) within uNK. (LFD: n=14; HFD: n=18; DIO: n=11; DIO-R: n=7). **C)** Representative MFI histograms and scatter plots showing expression levels of CD69 within NCR1+ uNK of LFD (n=14), HFD (n=18), DIO (n=11), and DIO-R (n=7) mice. Data represented as mean ± SD; *P*-values from unpaired Student’s t-tests for panels B&C.

### HFD impairs uterine spiral artery remodeling in mid but not late pregnancy

HFD-induced differences in uNK NCR1^+^ frequency and activity suggest that pregnancy associated processes controlled by uNK, like placental development^31^ and uterine artery remodeling^38^, may be affected by HFD. To assess this, we first examined if specific placental layers/zones at Gd14.5 are altered by HFD; Gd14.5 represents a developmental time-point when the mouse placenta is mature and fully functional. Total placental area (mm^2^), consisting of labyrinth zone (L), junctional zone (Jz), and decidua (D)^39^ were not different between control LFD and HFD mice (Figure 5A). Subgrouping of HFD placentas into DIO and DIO-R categories showed that diet-induced weight gain also did not affect overall placental area (Figure 5A). Moreover, HFD exposure did not result in changes in area within individual placental layers (i.e. L, JZ, and D) (Figure 5B, C, D). However, discrepancies in labyrinth area were identified between placentas of DIO and DIO-R mice, where the labyrinth area in DIO-R mice was greater (Figure 5B). Together our findings indicate that HFD-related uNK changes in early-mid gestation do not translate into major changes in placental morphology.

**Figure 5.**
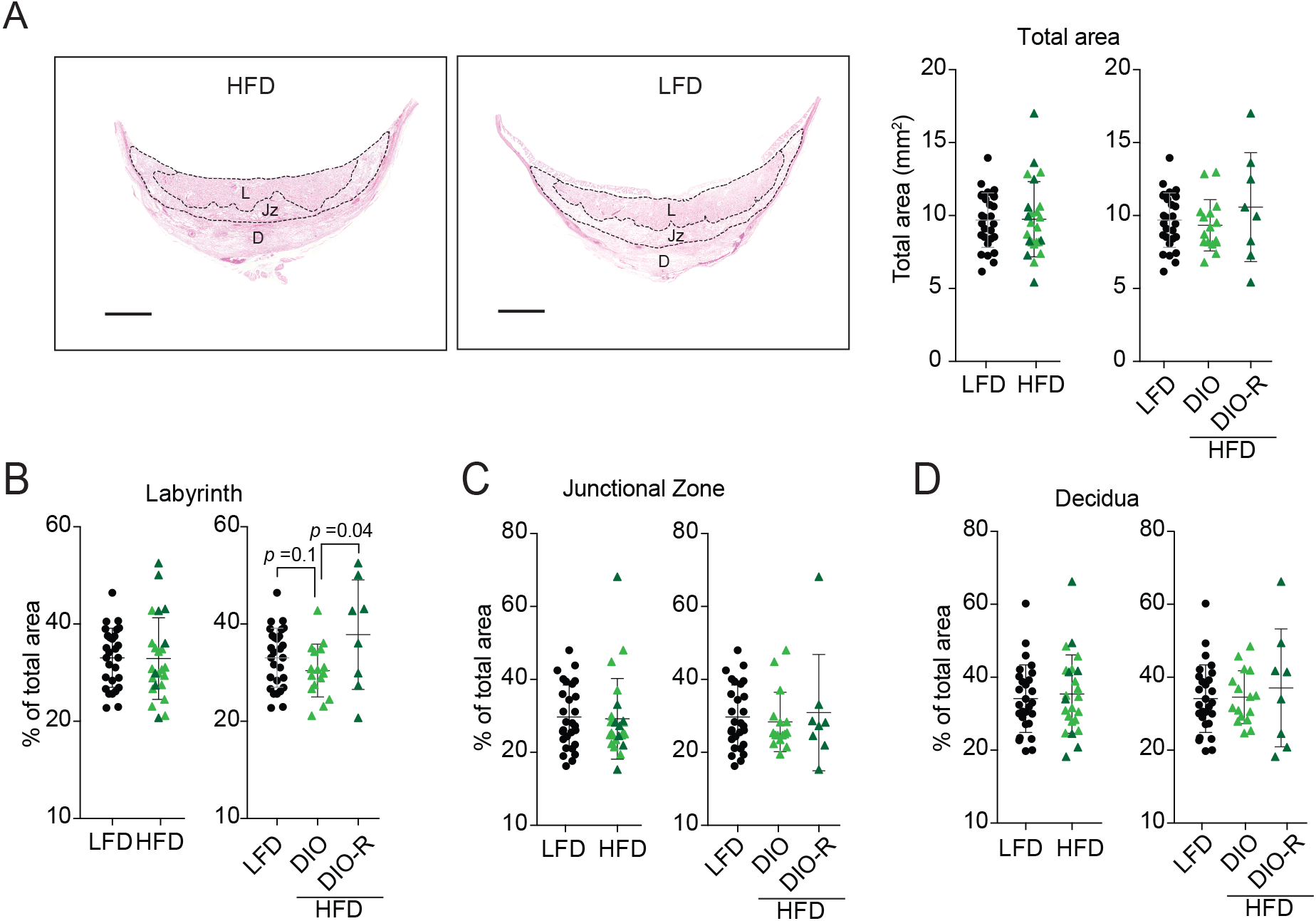
Exposure to HFD does not affect placental structure. **A)** Representative images of H&E-stained placentas from HFD- and LFD-fed mice with outlined Labyrinth (L), Junctional zone (Jz) and decidua (D). Scatter plots to the right show total placental area (mm^2^) from LFD, HFD, DIO and DIOR groups. Scale bar = 1000μm. Scatter plots showing proportional measurements of **B)** labyrinth, **C)** junctional zone and **D)** decidual areas relative to total placental area in LFD (n=28), HFD, (n=24), DIO (n=16), and DIO-R (n=8) mice. Data represented as mean ± SD; *P*-values from unpaired Student’s t-tests for panels A-D.

To examine whether HFD-induced uNK changes translate into alterations in uterine artery remodeling, measurements of Gd10.5 uterine arterial wall and vessel lumen areas, as well as their ratios, were next determined (Figure 6A). We also assessed decidual artery smooth muscle actin (SMA) expression, localization, and vessel intactness via immunofluorescence microscopy at Gd7.5, just prior to initiation of vascular remodeling, and at Gd10.5, when remodeling is at its peak and can be assessed by smooth muscle integrity by staining for SMA^40,41^. At Gd10.5, HFD-exposed mice had smaller vessels and thicker arterial walls (Figure 6A), and accordingly had significantly higher wall:lumen area ratios (Figure 6B). Interestingly, both DIO and DIO-R mice showed this arterial remodeling phenotype (Figure 6B). Examining SMA intactness at Gd7.5 by immunofluorescence microscopy, control LFD mice showed a marginal but significant increase in artery SMA coverage over HFD mice (Figure 6C, D). However, by Gd10.5, SMA staining within control mice was more sparse (i.e. smaller SMA:Perimeter ratio) than in HFD mice; HFD arteries showed only a partial loss in SMA staining, and this was reflected with higher SMA:Perimeter ratios (Figure 6D). By Gd14.5, differences in blood vessel remodeling observed within HFD mice were no longer observed (Figure 6E). Specifically, uterine arteries in both LFD and HFD mice showed dilated lumens and thin vessel walls indicative of proper remodeling (Figure 6E). Together, these findings suggest that maternal obesity disrupts uNK-mediated vascular remodeling in early-mid gestation, but that compensatory processes facilitating proper artery remodeling by mid-late gestation may overcome these initial impairments.

**Figure 6.**
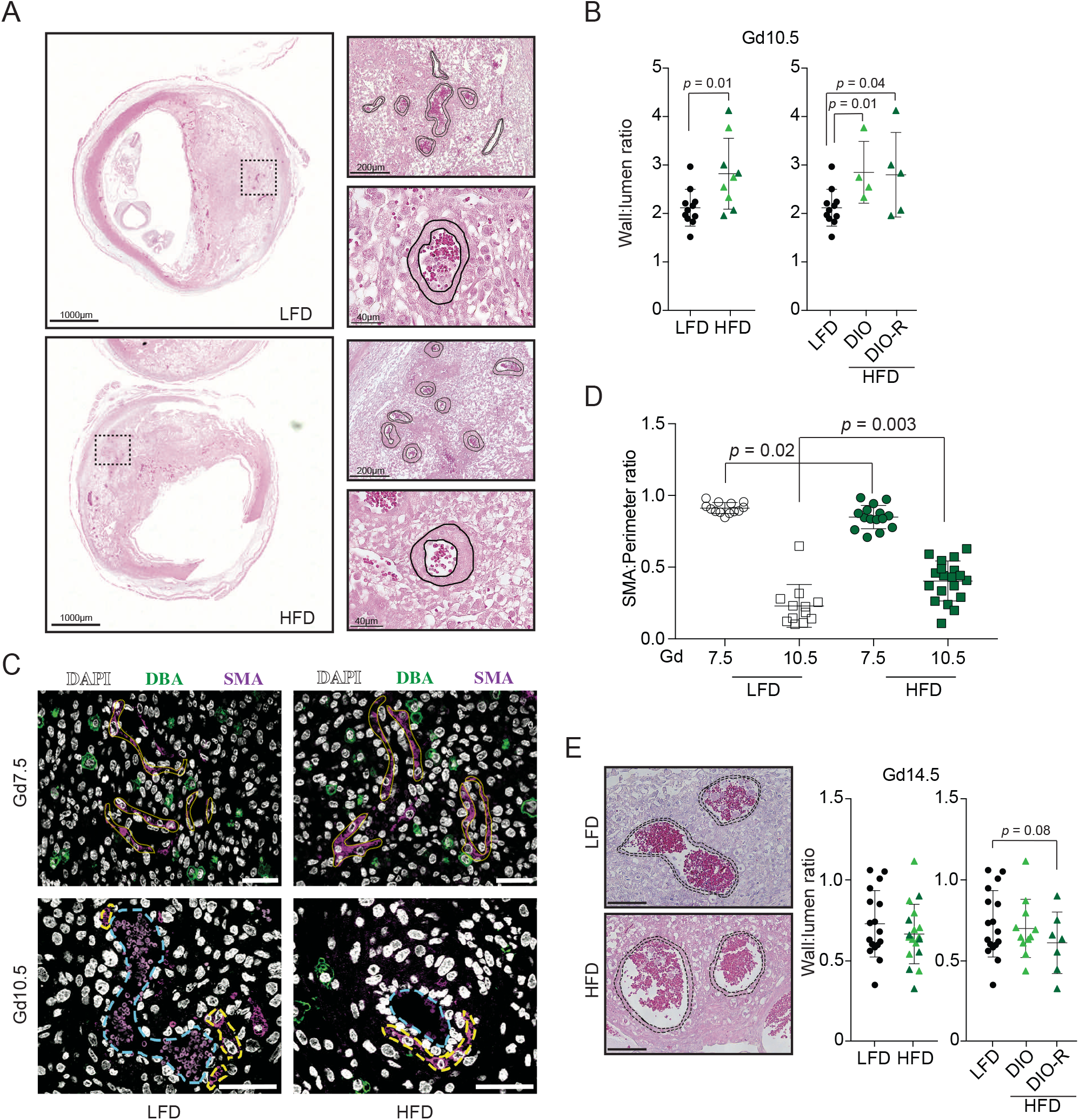
HFD leads to impaired spiral artery remodeling in mid pregnancy. **A)** Images of H&E stained Gd10.5 implantation sites. Hashed boxes and outlines show locations of spiral arteries being measured and outline vessel wall and vessel lumen areas. **B)** Arterial wall area (μm^2^) to lumen area (μm^2^) [wall:lumen] ratios from Gd10.5 LFD (n=11), HFD (n=9), DIO (n=4), and DIO-R (n=5) mice. **C)** Representative immunofluorescence images of Gd7.5 and Gd10.5 implantation sites stained with antibodies directed against smooth muscle actin (SMA; purple), and DBA (green). Nuclei are stained with DAPI (white). SMA surrounding arteries is highlighted with yellow dashed-lines, and artery lumens are indicated with blue-dashed lines. Scale bar = 40μm. **D)** Arterial SMA layer to arterial lumen ratios from Gd7.5 LFD (n=13) and HFD (n=15) mice, and in Gd10.5 LFD (n=12) and HFD (n=19) mice **E)** H&E images of GD14.5 decidual arteries in LFD (n=16), HFD (n=17), DIO (n=11), and DIO-R (n=6) mice. Scale bar = 100μm. Data represented as mean ± SD; *P*-values from unpaired Student’s t-tests for panels B, D&E.

### HFD exposure leads to subtle changes in uNK gene expression

To gain insight into possible uNK-related mechanisms altered by HFD exposure, global gene expression analyses were performed on mRNAs isolated from flow cytometry-sorted CD45^+^/CD122^+^/CD3^-^ uNK using gene microarrays (Figure 7A). Following standard probe filtering, normalization, and batch correction (Supplemental Figure 2A), differential gene expression comparison (FDR < 0.05), principal component (PCA), and hierarchical cluster analyses were performed between control LFD, DIO, and DIO-R uNK subgroups. Surprisingly, differential gene expression analysis did not identify any differentially expressed genes between the uNK subgroups (Supplemental Figure 2B), likely owing to the underpowered and heterogeneous composition of HFD uNK. PCA clustering based on global gene signatures further demonstrated that mouse uNKs exposed to HFD do not harbor unique or drastic differences in gene transcription, where uNK sub-group clustering showed DIO and DIO-R uNK to cluster randomly (Supplemental Figure 2C).

**Figure 7.**
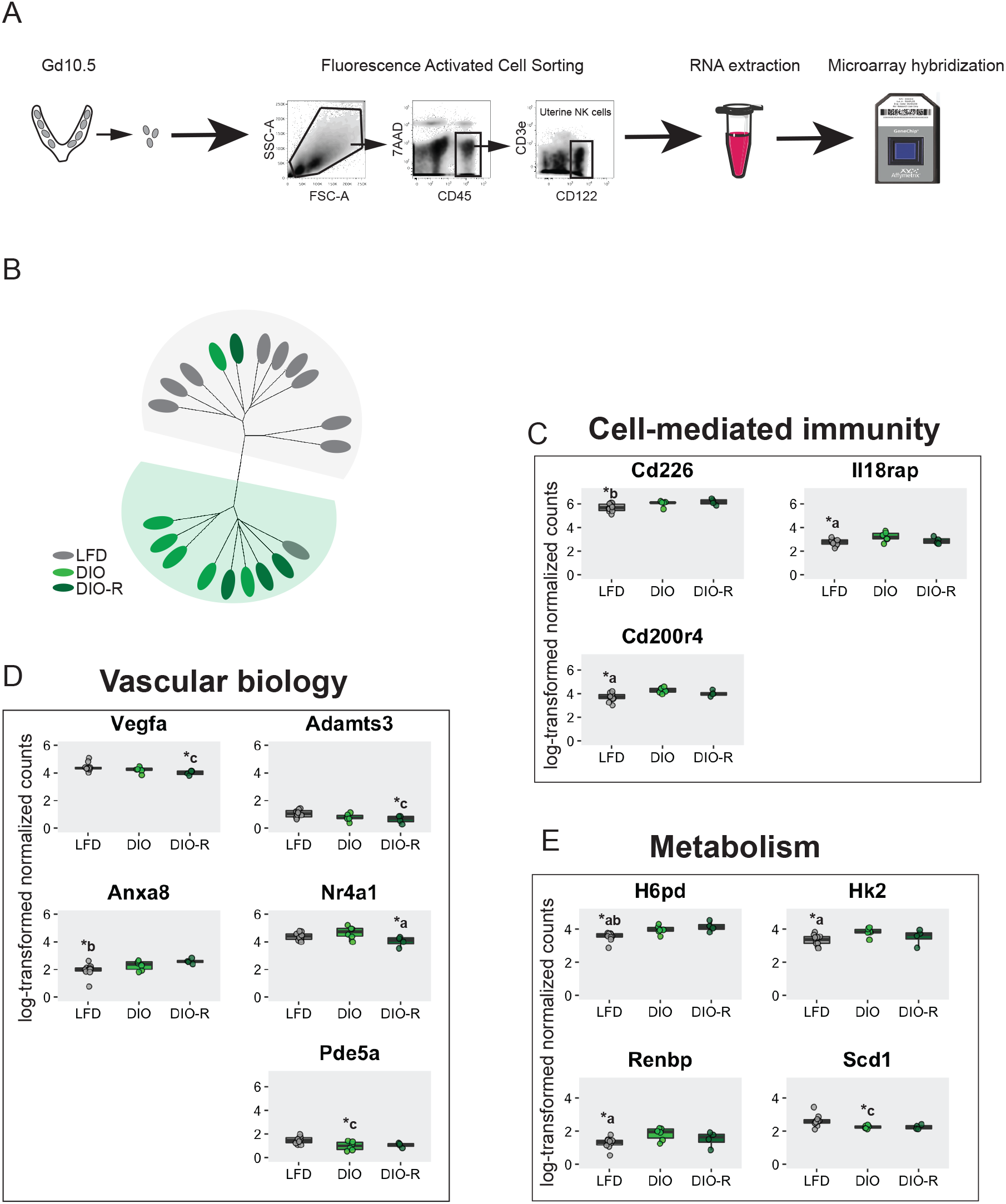
uNK gene expression profile of DIO and DIO-R vs LFD at Gd10.5. **A)** Schematic diagram depicting the workflow for gene expression analysis. uNK RNA was isolated from Gd10.5 implantation sites and hybridized onto Affymetrix microarray chip followed by analysis in R. **B)** Hierarchical clustering of LFD, DIO, and DIO-R uNK using a gene signature derived from nominal *P*-value ranking (*P*<0.05, fold-change > 1.25). Boxplots showing log-transformed, normalized expression levels for selected genes involved in **C)** immune function, **D)** vascular remodeling/angiogenesis and **E)** metabolism.

Despite the inability of a global gene signature approach to identify HFD-specific genes altered in uNK, hierarchical clustering of LFD, DIO, and DIO-R uNK using a 578 gene signature derived from raw *P*-value ranked genes (*P*<0.05, fold-change > 1.25) segregated uNK into diet-associated subgroups with 85-90% accuracy (Figure 7B). This finding suggests that HFD and associated weight gain may subtly affect the expression of specific uNK genes that were previously not identifiable due to the statistical stringency applied to the global high dimensional dataset. Of the selected 578 rank-ordered genes, one-way ANOVA with post-hoc Tukey HSD identified 71 genes to be differentially expressed within LFD, DIO, and DIO-R subgroups (Supplemental Table 1). Specifically, sets of genes linked with uNK cytotoxicity and activation were expressed at higher levels within both DIO (*Cd200r4, Ill8rap*) and DIO-R (*Cd226*) subgroups (Figure 7C). In line with our findings that uterine vascular remodeling was affected by high-fat diet, expression of pro-angiogenic/pro-remodeling *Vegfa, Adamts3, Nr4a1* and *Pde5a* were shown to be altered in DIO and DIO-R subgroups compared to LFD controls, with Vegfa-regulating *Anxa8* being inversely expressed within HFD samples compared to controls (Figure 7D). Targeted qPCR analysis of *Vegfa* in a subset of the same uNK mRNA samples used in the gene array supported the array’s finding (Supplemental Figure 2D). Moreover, HFD also resulted in alterations in levels of genes important in metabolism, where *H6pd* and *Hk2*, genes central to glucose metabolism, and *Renbp*, regulator of amino-sugar metabolism, were up-regulated in DIO and DIO-R subgroups (Figure 7E). Levels of *Scd1*, a gene important in lipid metabolism, were higher in the LFD control group (Figure 7E). Taken together, while chronic HFD exposure does not lead to robust global changes in uNK gene signatures, HFD does modestly affect the expression of select genes important in regulating uNK-mediated cytotoxicity/activation, vascular biology, and metabolism. These weight-gain dependent and independent differences provide insight into the biological underpinnings of how maternal obesity affects uNK processes important in pregnancy.

## DISCUSSION

In this study, we examine the effects of maternal HFD on uNK cell biology and uterine vascular transformations in early and mid-pregnancy. We demonstrate that HFD does not alter total uNK numbers or subsets of tissue-resident or conventional cells. However, weight gain-specific effects on truNK were observed, and additionally exposure to HFD results in a selective increase in NCR1^+^ uNK frequencies and overall uNK activity. These diet-related uNK changes associate with impairments in uterine artery remodeling in early-mid pregnancy, but were no longer observed by mid-late pregnancy, suggesting that a compensatory mechanism in uNK function may be taking place. While examination of global gene expression signatures did not identify specific uNK genes or pathways altered by HFD, gene signature-informed hierarchical clustering did segregate uNK according to diet exposure with relative high accuracy, indicating that uNK genes and gene pathways are moderately affected in obesogenic mice. Genes identified as being differentially expressed between control and HFD subgroups signified that uNK exposed to HFD harbor pro-cytotoxic and anti-angiogenic characteristics that together provide new insight into how maternal obesity may alter uNK processes critical to early pregnancy establishment and health.

To our knowledge, this is the first study to examine the effects of HFD on uNK cell proportions and states of activation in pregnancy. Importantly, this work also examines the specific effects of HFD-exposure and HFD-associated weight gain. While a previous study did report on the effects of maternal Western diet exposure on NK properties within draining inguinal lymph nodes of the uterus^10^, our study directly examines uNK tissue-resident and conventional subsets within the decidual mucosa. Because uNK are distinct from peripheral blood and lymphatic NKs^34,42,43^, our findings generate new insight into how maternal obesity potentially impacts immune cell function within the maternal-fetal interface. While HFD exposure does not affect the frequency of total leukocytes or uNK subtypes within the uterus, notably we show that HFD exposure increases both the proportion of NCR1^+^ uNK and uNK activation. Strikingly, differences in NCR1 frequency and uNK activation were primarily observed in HFD mice that were resistant to diet-induced weight gain (i.e. DIO-R), suggesting that the HFD-related features altered in uNK and uterine arteries may not necessarily be related to changes in overall adiposity. Our finding that weight-gain related effects on elevated blood serum TNFα and increased rates of fetal resorption may therefore be unrelated to changes in uNK biology. However, due to inter-sub-group heterogeneity and overall modest sample size, further work is needed to investigate the direct role of excessive adipose tissue (i.e. obesity) and diet-related factors like fatty acids and metabolites that may also lead to uterine immune cell dysfunctions.

The impact of obesogenic-diets on fetal health and pregnancy outcome in rodents has not been consistently reported on. Prior studies have reported that HFD exposure in pregnancy results in both reduced^44,45^ and increased^46,47^ fetal weight, and reduced placental labyrinth^48^ and junctional zone^44^ thickness, or no differences in placental weights^45,49,50^. In our study, placental and fetal weights and placental/decidual areas were not affected by HFD. Notably, HFD exposure did result in increased rates in fetal resorption, although this effect was modest. Important and somewhat counter to our findings showing that DIO-R uNK have elevated activity and expression of NCR1, the increase in fetal resorption is primarily attributed to DIO mice, and not weight gain-resistant DIO-R mice. The underlying cause for this observation is unknown but does nonetheless highlight the importance of needing to subcategorize mice subjected to experimental diet models of obesity as true weight gainers or as weight gain-resistant mice. In our hands, DIO and DIO-R murine uNK showed modest, but significant differences in specific parameters, and these differences were in part associated by weight gain-specific differences in gene expression.

Our focus on NCR1 stems from previous work reporting that *Ncr1^-/-^* mice exhibit defects in decidual angiogenesis and uterine vascular remodeling^36^. Notably, inhibition of NCR1 delays uNK cell maturation and activation, reflected by smaller cell diameters and reductions in cytokine production^51^. Our finding that HFD led to inverse associations between frequencies of NCR1^+^ uNK and degree of vascular remodeling (i.e. elevated NCR1 frequencies and impaired arterial remodeling) is inconsistent with what is known about the role of NCR1 in uNK biology. At least two possible explanations can account for this discrepancy. Firstly, the increase in NCR1^+^ uNK may reflect a level of compensation, and this is supported by the fact that initial impairments in vascular structure are no longer observed by Gd14.5.

A second possibility centers on the role of NCR1 as a key activating receptor important in promoting cytotoxic functions of NK^52–54^. Perhaps, elevated NCR1 frequency indicates that uNK exposed to HFD are either overly activated or are aberrantly cytotoxic. In line with this speculation, we show that mRNA levels of the pro-cytotoxic receptor *Cd226* (encoding for DNAM-1) are elevated within HFD uNK. DNAM-1 associates with NK maturation and memory and CD226^+^ NK are more cytotoxic and produce higher levels of inflammatory cytokines than do CD226^-^ counterparts^55,56^. Intriguingly, *Cd226* levels are more abundant within DIO-R, while transcripts encoding for CD200R4, a DAP12-associated activating receptor^57^, are expressed at higher levels in DIO uNK, suggesting that distinct mechanisms of activation are induced in relation to diet and/or adiposity/weight gain. Overall, these possibilities are consistent with a recent finding in human uNK that correlates maternal obesity to elevated rates of uNK degranulation^27^. However, it remains to be tested if obesity-linked hyper-activation in uNK equates to increased killing and dysfunction of cells within the maternal-fetal interface. Further, if indeed the case, cell-type specific uNK targets need also to be identified, as this will help in our understanding of the biological consequences of heightened uNK activity in response to maternal obesity and/or HFD.

Of interest, and consistent with previous findings in uNK from obese women^27^, HFD exposure in mice results in a decrease in *Vegfa* expression. While multiple cell types within the utero-placental environment are responsible for VEGF-A production, including decidual stromal cells^58,59^ and macrophages^60^, the transient nature of uNK in pregnancy and their links to vascular growth, branching, and remodeling suggest that uNK deficiency in VEGF-A production could directly translate into impairments in uterine vascular biology. VEGF-A is also a ligand for Flt-1 expressing trophoblasts, where VEGF-A plays roles in controlling trophoblast survival and proliferation^61–63^, however our finding that HFD exposure does not impact the development of specific zones of the placenta indicates that VEGF-A dysregulation may play a more significant and specific role in uterine artery angiogenesis.

It is becoming widely appreciated that cellular metabolism plays a role in regulating NK cell function and education^64^. A recent study demonstrated that glucose metabolism is required for receptor-activated NK cell IFNγ production^65^. Accordingly, Michelet *et al*^66^ showed that obesity-driven metabolic paralysis results blunted NK cell cytotoxic responses. In line with these findings we demonstrate an increase in expression of *H6pd* and *Hk2* and decreased levels of *Scd1* in DIO and DIOR uNKs, where *H6pd* and *Hk2* are important regulators of glycolysis and *Scd1* is involved in fatty acid metabolism. Overall, it is likely that the obesogenic environment in HFD mice may induce a metabolic switch in uNKs that impacts maturation and function that may ultimately alter maternal-fetal interface remodeling and overall uNK education. Potential metabolic effects on uNK processes will be of high interest to further examine as metabolic dysfunction and altered programming of uNK in human pregnancies could in part contribute to impaired pregnancy outcomes.

uNKs are specialized immune cells that play an important role in coordinating reproductive success. Deviations in normal uNK function may contribute to defective vascular modifications or aberrant induction of cytotoxic potential at the maternal-fetal interface. Such uNK-related changes may play roles in pregnancy complications. Our study provides insight into how exposure to an obesogenic diet alters uNK activity and alters spiral artery remodeling, highlighting possible cellular mechanisms underlying HFD exposure. Maternal obesity continues to pose serious consequences on reproductive health, thus future studies aimed to further dissect how obesity-driven immune and cellular alterations impact pregnancy health will be of substantial importance.

## METHODS

### Animals

Mouse experiments were carried out in accordance with the guidelines of the Canadian Council on Animal Care and approved by SFU University Animal Care Committee (protocol 1094) and the UBC Animal Care Committee (protocol A17-0060). C57BL/6J mice were purchased from the Jackson Laboratory (Bar Harbor, ME, USA). Animals were housed under standard conditions in groups of max. 5 mice in individually ventilated cages with water and food available *ad libitum*, under a 12:12h light/dark cycle.

### High-fat/high sugar obesogenic model

At 9 weeks of age, females were placed on either a high-fat, high-sucrose diet (HFHS; 45% kcal fat, 35% carbohydrate (including 17% kcal sucrose), 20% kcal protein, 4.73 kcal/g, D12451, Research Diets, New Brunswick, NJ) or a nutrient-matched low-fat, no-sucrose control diet (CON; 10% kcal fat, 70% kcal carbohydrate (corn starch and maltodextrin), 20% kcal protein, 3.85 kcal/g, D12450K, Research Diets). Females were kept on the experimental diets for 13 weeks prior to being paired with male studs. The appearance of a plug the next morning was taken as day 0.5 of pregnancy. Because not all females became pregnant when first paired, some had been on the experimental diet for 13–19 weeks at the start of pregnancy. On days 7.5, 10.5 and 14.5 of pregnancy females were euthanized and their blood collected via cardiac puncture. The utero-placental tissues with multiple implantation sites were collected from each mouse.

### ELISA measurements of serum TNFα

Serum was purified from whole blood of females sacrificed at Gd10.5 of pregnancy using serum separator tubes following standard procedures. Serum levels of TNFα were determined using commercially available Mouse TNFα ELISA kit (R&D systems) according to manufacturer’s protocol.

### Decidual leukocyte preparations

For examination of uNK cell populations, murine implantation sites were separated from uterine tissues and were mechanically and enzymatically processed to obtain single-cell suspensions. In brief, tissues were washed extensively in cold phosphate-buffered saline (PBS; pH 7.4), and following this, tissues were minced with a razor blade and subjected to a 30-minute enzymatic digestion at 37 °C in 3 mL of 1:1 DMEM/F12 media (Gibco, Grand Island, NY; 200 mM L-glutamine) with 1X-collagenase/hyaluronidase (10X stock; StemCell Technologies, Vancouver, Canada), 80 μg/mL DNaseI (Sigma, St. Louis, MO), penicillin/streptomycin, and Anti antimycotic solution (100X dilution, Gibco). Single cell suspensions were passed through a 100μm strainer and decidual leukocyte enrichment was performed by Percoll (GE Healthcare) density gradient (layered 40%/80%) centrifugation. To avoid red blood cell contamination, cell mixture was briefly incubated with NH_4_Cl. Decidual leukocytes were immediately used for uNK isolation by fluorescence activated cell sorting (FACS) or cell-surface marker characterization via flow cytometry analyses.

### Flow cytometry

Flow cytometry analyses were carried out following a standard protocol. In brief, 0.5×100צ isolated leukocytes were incubated with Fc Block (anti-CD16/CD32 antibody at 1:35 dilution in PBS; BD Pharmingen) for 10 min on ice to prevent non-specific binding. Cell suspensions were further incubated with fluorescent-conjugated antibodies directed against cell-surface markers for 30 minutes at 4°C. The following antibodies were used: APC ef780 CD3e (1:50; clone 145-2C11; eBioscience), ef450 CD45 (1:100; clone 30-F11; eBioscience), PerCP710 CD122 (1:100; clone TM-b1; eBioscience), APC CD122 (1:35; clone TM-Beta 1; BD Pharmingen), APC CD49a (1:50; clone HMα1; BioLegend), PeCy7 CD49b (1:100; clone DX5; eBioscience), APC CD69 (1:25; clone H1.2F3; eBioscience), PeCy7 NKp46 (1:50; clone 29A1.4; eBioscience). Following staining, cells were resuspended in 7-AAD viability staining solution (at 1:25 dilution in PBS+ 2%FBS +1% penicillin/streptomycin; eBioscience). Samples were acquired on an LSRII FACS (BD Biosciences) and data was analysed by FlowJo 5.0 software (Tree Star, Inc., Ashland, OR, USA).

### Immunohistochemistry, IF microscopy and morphometry

Implantation sites from Gd7.5, Gd10.5 and Gd14.5were fixed in 4% paraformaldehyde for 24-48h followed by processing into paraffin blocks. Six-μm sections were stained with hematoxylin and eosin (H&E) according to standard methods. For IF staining, tissues underwent antigen retrieval by heating slides in a microwave for 5 minutes in 30-second intervals in a sodium citrate buffer (pH 6.0). Sections were then incubated in a blocking solution (5% bovine serum albumin (BSA) in Tris-buffered saline with 0.05% Tween-20 (TBST)) for 1 hour at room temperature. Further, samples were incubated with the following antibodies overnight at 4°C: rabbit polyclonal alpha-smooth muscle actin (1:200; Abcam) and biotinylated DBA (1:100; Vector Laboratories). Slides were incubated with Alexa Fluor 568/488–conjugated streptavidin and goat anti-rabbit secondary antibodies (Life Technologies) for 1 hour at room temperature. Glass coverslips were mounted onto slides using ProLong Gold anti-fade reagent containing DAPI (Life Technologies).

#### DBA^+^ cell assessments

DBA^+^ cell quantities in Gd7.5 and Gd10.5 implantation sites were calculated based on DBA fluorescence intensity threshold. Slides were imaged using a 20x Plan-Apochromat/0.80 NA or a 40x EC-Plan-Neofluar/0.9 PoI objective (Carl Zeiss). Data was processed and analyzed using ZenPro software (Carl Zeiss). DBA^+^ cell diameters in Gd10.5 implantation sites were measured by randomly selecting 100 DBA^+^ uNK cells from each implantation site. Slides were imaged using a 40x EC-Plan-Neofluar/0.9 PoI objective (Carl Zeiss). Using ImageJ v1.50 software (NIH), the diameters of these cells were measured across their longest and shortest axes; an average of both values was used for statistical analysis. All images were obtained using Axiocam 506 monochrome digital camera (Carl Zeiss).

#### Spiral artery remodeling and SMA layer assessments

Spiral artery remodeling measurements were performed on H&E stained slides. For Gd10.5, (HFD: n=9 implantation sites from 8 independent pregnancies; LFD: n=11 implantation sites from 9 independent pregnancies) all of the spiral arteries within the decidua were measured in triplicate (three sections 48μm apart from each other). For Gd14.5, (HFD: n=18 implantation sites from 7 independent pregnancies; LFD: n=17 implantation sites from 6 independent pregnancies) all of the spiral arteries within the decidua were measured in duplicate (two sections 48μm apart from each other). Size of arterial walls and lumens was determined by measuring cross-sectional areas with ImageJ v1.50 software (NIH). Wall:lumen ratios were calculated by dividing the arterial wall area (μm^2^) into the artery lumen area (μm^2^). Slides were imaged using a 20x Plan-Apochromat/0.80 NA objective (Carl Zeiss). All images were obtained using Axiocam 105 color digital camera (Carl Zeiss). SMA layer intactness was measured in spiral arteries of Gd7.5 (HFD: n=15 arteries from 4 independent pregnancies; LFD: n=13 arteries from 4 independent pregnancies) and Gd10.5 (HFD: n=19 arteries from 8 independent pregnancies; LFD: n=12 arteries from 7 independent pregnancies). Slides were imaged using a 40x EC-Plan-Neofluar/0.9 PoI objective (Carl Zeiss). Arterial length and SMA layer were measured using ImageJ v1.50 software (NIH). SMA:Perimeter ratios were calculated by dividing SMA^+^ vascular perimeter (μm) over total arterial perimeter (μm). All images were obtained using Axiocam 506 monochrome digital camera (Carl Zeiss).

#### Placental morphology assessments

H&E stained slides of Gd14.5 implantation sites were used for placental morphology measurements. Total area (mm^2^) and areas of each individual placental layer (mm^2^) were measured using ImageJ v1.50 software (NIH). To normalize for differences in total placental area among the measurements, individual layer area was expressed as a percentage of total area at each measurement point. Slides were imaged using 10x N-Achroplan/0.25 Ph1 objective (Carl Zeiss). All images were obtained using Axiocam 105 color digital camera (Carl Zeiss).

### RNA extraction

Prior to RNA extraction, CD45+CD3-CD122+ uNK cells from Gd10.5 implantation sites were isolated by FACS using a FACSAria flow cytometry (BD Biosciences). Total RNA was prepared from isolated uNKs using TRIzol LS reagent (Ambion) following by RNeasy MinElute Cleanup (Qiagen) and DNase treatment (Ambion) according to manufacturers’ protocols. RNA purity was confirmed using a NanoDrop Spectrophotometer (Thermo Fisher Scientific) and Agilent 2100 Bioanalyzer (Agilent). Only RNA samples having an RNA integrity number (RIN) greater than 8.0 were used.

### Microarray analysis

Total RNA samples extracted from mouse uNKs were sent to Génome Québec Innovation Centre (McGill University, Montréal, Canada) for RNA quantification. Briefly, mouse RNA samples were prepared for transcriptome profiling using the GeneChip™ Pico Reagent Kit (Affymetrix) as per manufacturer’s protocol. Samples were run on the Clariom™ S Mouse Array to measure gene expression at >20000 genes in the mouse genome. Raw data generated from the arrays were read into R statistical software (version 3.5.1) with the Bioconductor *oligo*^67^ package to convert raw Affymetrix CEL files into an expression matrix of intensity values. Within oligo package samples were background corrected, normalized, and log-transformed. Probe filtering identified 718 probes that mapped to multiple gene symbols were excluded from the dataset. Additionally, 6923 Affymetrix control probes and 749 probes that were not annotated to any given gene were also removed. A “soft” intensity-based probe filtering^68^, as recommended by *limma*, was performed to eliminate 2181 low variance and low intensity probes. Together, these measures eliminated 10571 probes, leaving a total of 18558 probes remaining for further analysis. Using ComBat from *sva* package, known technical variation associated with batch was removed. Following preprocessing, differential expression analysis between groups was performed by applying a linear model to expression values using the *limma* package’s moderated t-statistics with empirical Bayesian variance estimation^69^.

### cDNA synthesis, and qPCR

RNA was reverse-transcribed using a first-strand cDNA synthesis kit (Quanta Biosciences) and subjected to qPCR (ΔΔCT) analysis, using PerfeCTa SYBR Green FastMix Low ROX (Quanta Biosciences) on an ABI ViiA 7 Real-Time PCR system (Thermo Fisher Scientific). Forward and reverse primer sets used as previously described: *VEGFA* (F: 5’-GTGCACTGGACCCTGGCTTTA-3’, R: 5’-GGTCTCAATCGGACGGCAGTA-3’)^70^; *HPRT* (F: 5’-CTGGTGAAAAGGACCTCTCG-3’, R: 5’-TGAAGTACTCATTATAGTCAAGGGCA-3’)^71^. All raw data were analyzed using QuantStudio V1.3 software (Thermo Fisher Scientific). The threshold cycle (CT) values were used to calculate relative gene expression levels. Values were normalized to *HPRT* transcripts.

### Statistical analysis

Unless stated otherwise, statistical significance for all data was assessed by unpaired two-tailed Student’s t-tests. *P* <0.05 was considered significant. Statistical analyses were performed using Prism7 software (GraphPad Software, Inc.).

## ACKNOWLEDGEMENTS

We are grateful to staff from Simon Fraser University Animal Care facility for their assistance. We would like to extend our gratitude to BCCHRI Flow Core Facility Manager, Dr. Lisa Xu, for her expertise and assistance in cell sorting of uNK cells. We are grateful to staff at Génome Québec Innovation Centre at McGill University for their help with microarray hybridization.

## AUTHOR CONTRIBUTIONS

AGB and JKC designed the research. JB, CK, BC, DM, JKC and AGB performed the experiments. JB, CK, WPR, AGB and JKC analysed the data and wrote the paper. All authors read and approved the manuscript.

## SUPPLEMENTAL FIGURE LEGENDS

**Supplemental Figure 1.** Scatter plots of **A)** placental and **B)** fetal weights at Gd14.5. (LFD: n=17; HFD: n=16; DIO: n=10; DIO-R: n=6). **C)** DBA+ cell diameter measurements (μm) and photomicrographs of DBA (DBA: red, DAPI/nuclei: blue) immunofluorescence staining showing the approach for cell diameter measurements (yellow dashed lines). Scale bar = 10μm. (LFD: n=9 i.s.; HFD: n=9 i.s.; DIO: n=4 i.s.; DIO-R: n=5 i.s.). Data represented as mean ± SD; *P*-values from unpaired Student’s t-test for panels A-C.

**Supplemental Figure 2.** uNK gene expression analysis. **A)** Principal component analysis (PCA) of raw data, normalized data, filtered data and ComBat batch corrected data. **B)** Volcano plots following differential gene expression analysis of HFD vs LFD, DIO vs LFD and DIO-R vs LFD groups. **C)** PCA clustering based on global gene signatures in LFD, DIO and DIO-R groups. **D)** Scatter plot of relative mRNA expression of *Vegfa* in Gd10.5 uNK (HFD: n=5; LFD: n=5). Data represented as mean ± SD; *P*-values from unpaired Student’s t-test.

