## Supplementary figures and images for "Obesogenic diet exposure alters uterine natural killer cell biology and impairs vasculature remodeling in mice"

### Supplemental Figure 1

#### Supplemental Figure 1

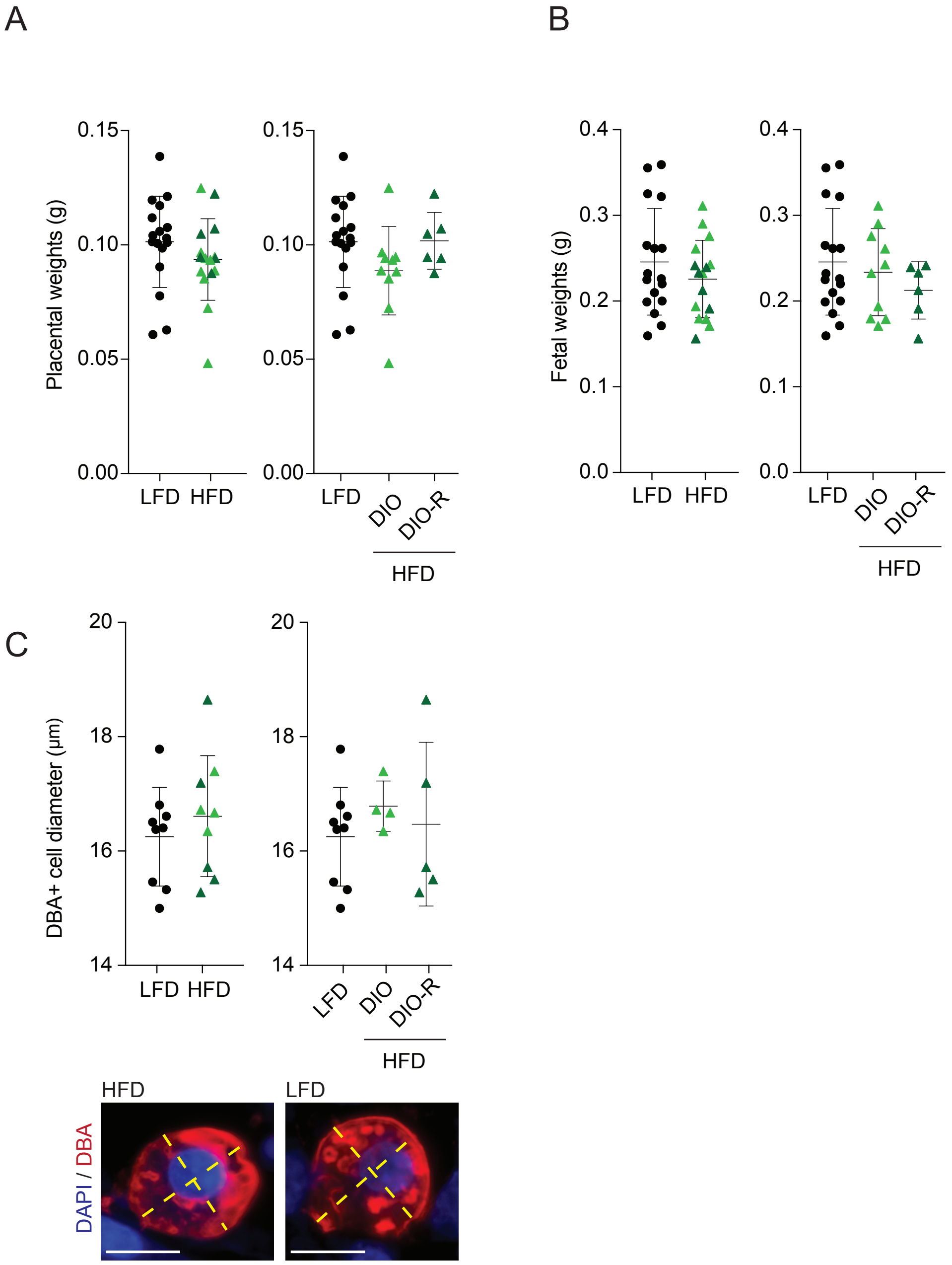

### Supplemental Figure 2

#### Supplemental Figure 2

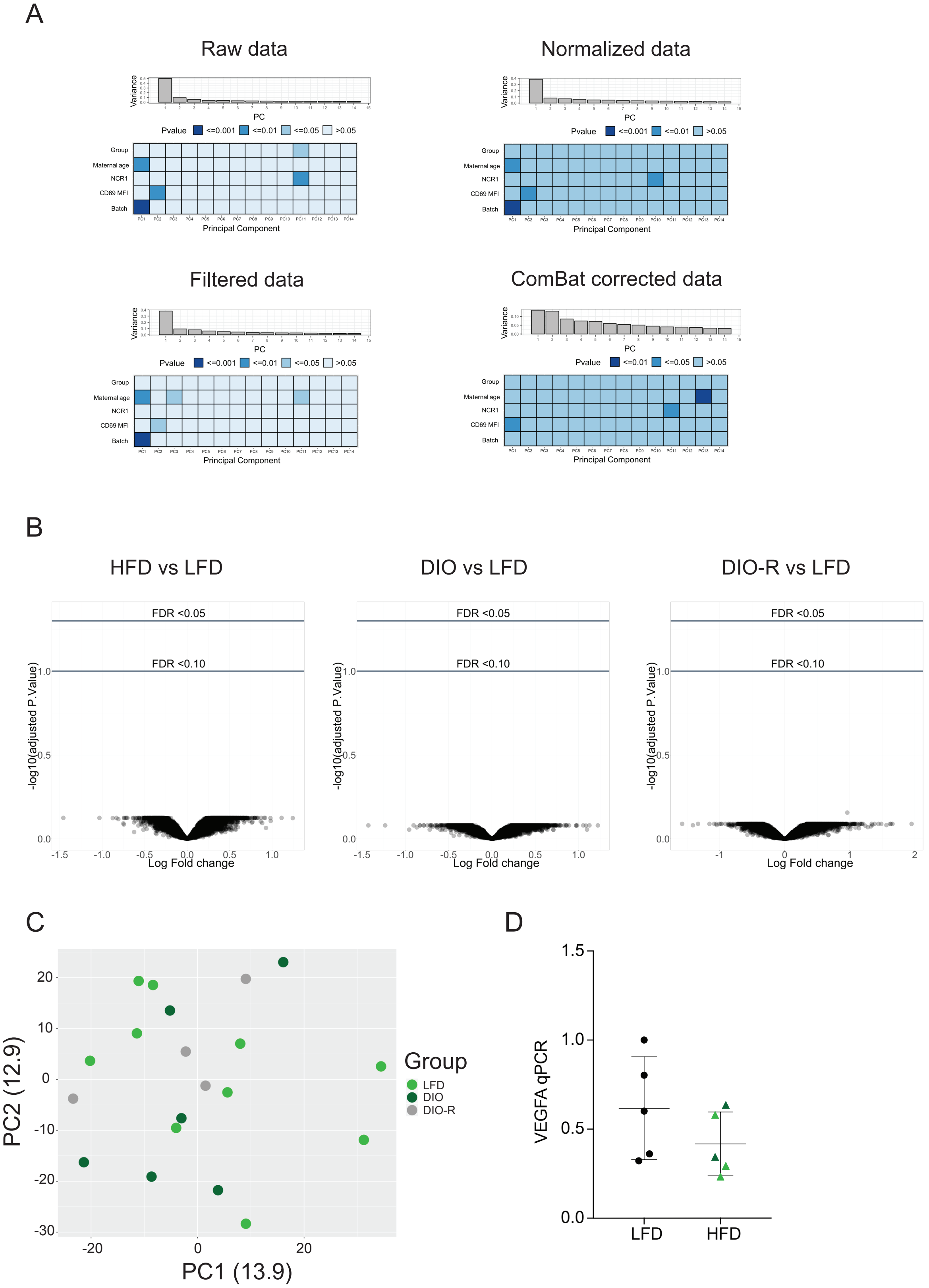
